# Intralumenal docking of Cx36 channels in the ER isolates mis-trafficked protein

**DOI:** 10.1101/2022.07.15.500247

**Authors:** Stephan Tetenborg, Viktoria Liss, Leonhard Breitsprecher, Ksenia Timonina, Anna Kotova, Alejandra Jesús Acevedo Harnecker, Chunxu Yuan, Eyad Shihabeddin, Karin Dedek, Georg Zoidl, Michael Hensel, John O’Brien

## Abstract

The intracellular domains of connexins are essential for the assembly of gap junctions. For connexin 36 (Cx36), the major neuronal connexin, it has been shown that a dysfunctional PDZ binding motif interferes with electrical synapse formation. However, it is still unknown how this motif coordinates the transport of Cx36. In the present study, we characterize a phenotype of Cx36 mutants that lack a functional PDZ binding motif using HEK293T cells as an expression system. We provide evidence that an intact PDZ binding motif is critical for proper ER export of Cx36. Removing the PDZ binding motif of Cx36 results in ER retention and the formation of multi-membrane vesicles containing gap junction-like connexin aggregates. Using a combination of site directed mutagenesis and electron micrographs we reveal that these vesicles consist of Cx36 channels that docked prematurely in the ER. Our data suggest a model in which ER-retained Cx36 channels reshape the ER membrane into concentric whorls that are released into the cytoplasm.

## Introduction

Electrical synapses serve as a fast means for signal transmission in the nervous system and provide unique functions such as neuronal synchronization or signal averaging (Bloomfield and Volgyi, 2009; Christie et al., 2005). Structurally they are defined as gap junctions, specialized cell junctions that contain clusters of intercellular channels establishing a conductive link for ionic currents and metabolites (Kumar and Gilula, 1996). Each gap junction consists of thousands of intercellular channels that are made of membrane proteins called connexins. A characteristic feature of these proteins is their ability to self-assemble into gap junction channels. To form such a channel, six connexins first oligomerize and assemble into a connexon (or hemichannel). Afterwards, two opposing connexons on adjacent cell membranes dock via non-covalent interactions of the extracellular loops in each connexin to form a functional gap junction channel. Among all gap junction proteins that have been described in mammals, connexin 36 (Cx36) is regarded as the major neuronal connexin because its expression is virtually (except for the pancreas (Serre-Beinier et al., 2000) confined to neurons (Condorelli et al., 1998). Gap junctions that are made of this particular isoform are characterized by their low single channel conductance and their tremendous degree of plasticity (O’Brien, 2014; Srinivas et al., 1999). Especially, neuromodulators and second messenger systems have been shown to exert drastic effects on gap junction coupling via phosphorylation of Cx36 (Kothmann et al., 2009; Kothmann et al., 2012). Recent studies indicate that active transport and turnover mechanisms of Cx36 function as an additional means to adjust the strength of electrical synapses (Brown et al., 2019; Flores et al., 2012).

Like many membrane proteins, connexins follow a characteristic life cycle. Depending on the isoform, they either oligomerize in the endoplasmic reticulum or the Golgi apparatus. From here, newly assembled channels are transported to the plasma membrane by cytoskeleton dependent mechanisms and added to the periphery of the gap junction. The internalization of gap junctions relies on the formation of so-called annular junctions (Bell et al., 2018; Piehl et al., 2007; Raviola et al., 1980). These structures are formed by the endocytosis of gap junction channels of both cells from the center of the plaque leading to the formation of a double membrane vesicle that is subjected to the proteolytic systems of the cells (Falk et al., 2016; Segretain and Falk, 2004). Significant insight into the life cycle of gap junction proteins was gained by imaging studies of recombinant connexins in expression systems. Usually, a GFP-tag (or any other fluorophore) is fused to the intracellular domains of a connexin to monitor transport processes or simply to visualize the protein without an immuno-labeling procedure (Bukauskas et al., 2000; Falk et al., 2016; Thomas et al., 2005). Protein tags, although they have proven to be powerful tools, can compromise protein function depending on their insertion site. Often, they act as a physical barrier and mask certain protein-protein interaction domains, resulting in a loss of association with important binding partners (Helbig et al., 2010). Such effects have been described for Cx43 and Cx36. The expression of a Cx43-GFP fusion protein in which a GFP tag is linked to the C-terminus of Cx43 leads to the formation of unusually large gap junctions in HeLa cells (Hunter et al., 2003). For Cx36-EGFP, on the contrary, a C-terminal GFP tag prevents gap junction formation in transfected HeLa cells and results in a partial loss of electrical synapses in transgenic mice (Helbig et al., 2010; Meyer et al., 2014). These phenotypes can be explained by the position of the GFP tag. In each of the constructs, GFP is located in immediate proximity to the PDZ binding motif, a short amino acid sequence at the C-terminus of connexins that is necessary to recruit scaffold proteins containing PDZ domains. The PDZ binding motif of Cx36 consists of the four C-terminal amino acids: SAYV: Serine (S), Alanine (A), Tyrosine (Y) and Valine (V) (Li et al., 2004). Due to its proximity to the GFP tag, this sequence is inaccessible for PDZ domains preventing protein-protein interaction at the C-terminus, which ultimately results in the transport deficit observed in Cx36-EGFP mice and transfected HeLa cells. These effects can be reproduced with Cx36 mutants that lack the PDZ binding motif. Although these mutants show a capacity to form gap junctions, they only assemble into intercellular clusters that are significantly smaller than those of wild type Cx36. This suggests that the transport deficit of Cx36-EGFP is based on a dysfunctional carboxyl terminus.

Accumulating evidence suggests that the PDZ binding motif of Cx36 is critical for electrical synapse formation. Still, it is unclear how this sequence coordinates the transport of Cx36. Interestingly, Helbig et al., (2010) reported that the Cx36-EGFP fusion protein not only fails to form gap junctions in HeLa cells, but it also seems to assemble into spherical cytoplasmic clusters. Similar observations were made for Cx36 mutants that lack the tubulin binding motif. Brown et al., (Brown et al., 2019) have shown that deleting this motif (ranging from W277 to S292) interrupts the microtubule-dependent transport of Cx36 and causes the formation of large annular gap junctions. Each of these two studies describe a transport deficit that correlates with the increased formation of intracellular vesicles. How these structures are formed and why they occur in Cx36 mutants with a dysfunctional carboxyl terminus is currently unknown.

In the present study we describe a transport deficit that is observed in PDZ binding deficient Cx36 mutants (throughout the paper referred to as Cx36/S318ter). We identify a mechanism that explains the formation of cytoplasmic connexin aggregates that have been reported in earlier studies and reveal how deleting the PDZ binding motif affects the intracellular transport of Cx36. Overexpression of the Cx36/318ter mutant results in the formation of multi membrane vesicles that originate from the ER. Our data suggest that deleting the PDZ binding motif prevents Cx36 from exiting the ER. Because of this retention mechanism, connexons on adjacent ER membranes are allowed to dock prematurely, leading to the formation of gap junction-like ER sheets that coil up into multi-membrane vesicles called connexin whorls. Here, we reveal an unanticipated function of the classical head-to-head docking mechanism of gap junction channels and show that it can function as a release mechanism to isolate ER retained connexins.

## Methods

### Cell Culture and Transient Transfection

Human embryonic kidney 293T cells (HEK293T/17; catalog #CRL-11268; ATCC, Manassas, VA, USA; RRID: CVCL_1926) were cultivated in Dulbecco’s Modified Eagle Medium (DMEM) supplemented with 10% fetal bovine serum (FBS), 1% penicillin and streptomycin, and 1% non-essential amino acids (all Thermo Fisher Scientific, Rockford, IL, USA) at 37°C in a humidified atmosphere with 5% CO2. For the cell surface biotinylation assay, ∼800,000 cells were seeded in 60 mm plates. For live cell imaging, ∼25,000 cells were seeded in 35 mm glass-bottom dishes (MatTek Corporation, Ashland, MA, USA). For live microscopy experiments, the medium was changed to DMEM lacking phenol red 30 min prior to imaging. HEK293T cells were transiently transfected with Effectene™ Transfection Reagent Kit (Qiagen Inc., Valencia, CA, USA) according to the manufacturer’s guidelines. Cells were transfected with a total of 1000 ng of DNA for each 60 mm plate. For 35 mm glass-bottom dishes, cells were transfected with 200 ng of DNA.All assays were performed 48 h post-transfection. For immunocytochemical experiments 270,000-450,000 cells were plated on to 35 mm dishes a day before transfections. 1 µg of DNA were transfected using 5 µl Geneporter2 (Genlantis) or 1.5 µl Lipofectamine 2000 (Thermo Fisher).

### DNA constructs

Cx36 S318ter, C55S and C62S mutants were generated using the Q5 site directed mutagenesis kit (New England Biolabs). The Cx36 pcDNA plasmid was PCR amplified with modified primers to introduce the corresponding deletions or substitutions. The following primers were used: S318ter: Forward: 5`TGA AAG GGC AGG TTT GGG GAA3`; Reverse: 5` GTC ACT GGA CTG AGT CCT GCC 3`; C55S: Forward: 5` ATG TTT GTG TCC AAC ACC CTA C 3`; Reverse: 5` GGT CTG CTC ATC ATC GTA CA 3`; C62S: Forward: 5` CAG CCC GGC TCT AAC CAG GC 3`; 5` TAG GGT GTT GCA CAC AAA CAT GG 3`. The Cx36-SNAP construct was cloned using Gibson assembly (New England Biolabs). The SNAP plasmid was obtained from Dr. Ana Egana (Addgene plasmid 101124) and used as a template to clone the coding sequence of the enzyme into the Cx36 pcDNA vector (C-terminal tag). The dGBP-TurboID construct was obtained from Dr. Thomas Halls` lab (Addgene plasmid 163857; kindly provided by Dr. Adam Miller (University of Oregon) before this construct was available on Addgene and cloned into the V5-TurboID-NES pcDNA Vector (Addgene plasmid 107169) to substitute TurboID with dGBP-TurboID. The ER marker GFP-SEC61B (Addgene plasmid 121159) and DsRed2 ER5 (Addgene plasmid 55836) originated from Dr. Christine Mayr and Dr. Michael Davidson, respectively.

### BioID pull-down

For large scale BioID experiments 4×10^6^ HEK293T cells were plated onto 100 mm dishes and co-transfected the next day with 4 µg Cx36-EGFP, 4 µg Cx36-SNAP and 4 µg V5-dGBP-TurboID using 50 µl Geneporter2 (Genlantis). 24 h after transfection HEK293T cells were treated with 50 µM Biotin in 10% DMEM for 3 h. After biotin supplementation, HEK293T cells were collected and rinsed in PBS. Cells were lysed in lysis buffer (1% Triton X-100, 50 mM Tris pH 7.5, 250 mM NaCl, 5 mM EDTA, 50 mM NaF, 1 mM Na_3_VO_4_, 1 mM DTT and protease inhibitors (Roche) and incubated on ice for 30 min. The lysates were centrifuged at 16,000g for 10 min. 10 mg of the supernatant were mixed with 275 µl of streptavidin beads myOne C1 (Thermofisher, Dynabeads MyOne Streptavidin C1) and incubated overnight at 4**°**C on a rotating platform. At the next day streptavidin beads were collected using a magnetic stand and washed in the following washing buffers (10 min for each washing step): 2x in wash buffer 1 (2% SDS), once with wash (0.1% deoxycholate, 1% Triton X-100, 500 mM NaCl, 1 mM EDTA and 50 mM Hepes, pH 7.5), once with wash buffer 3 (250 mM LiCl, 0.5% NP-40, 0.5% deoxycholate, 1 mM EDTA and 10 mM Tris, pH 8.1) and twice with wash buffer 4 (50 mM Tris, pH 7.4, and 50 mM NaCl). Proteins were eluted with 2 × 35 µl 1x Laemmli sample buffer (BioRad, 1610747) at 95**°**C.

### Pulse-chase experiments

For pulse-chase labeling experiments, Cx36-SNAP transfectants were treated with 6 µM SNAP-Cell Oregon Green (New England Biolabs Inc., Boston, MA, USA) in 10% FBS DMEM for 30 min at 37°C to quench all Cx36-SNAP molecules. Afterwards coverslips were washed 3 times in serum free DMEM and incubated for 30 min at 37**°**C. At different time points after the initial incubation (0.25 h, 0.5 h, 1 h, 2 h, 3 h) the cells were incubated in 3 µM SNAP-Cell TMR-Star (New England Biolabs Inc., Boston, MA, USA) in 10% FBS DMEM for 30 min and embedded using Vectashield (supplemented with DAPI).

### Cell Surface Biotinylation Assay

HEK293T cells were seeded on 60 mm plates and transfected with Cx36-WT-SNAP or Cx36-C62S-SNAP. Biotinylation assay was performed 48 h post-transfection. Cells were washed once with PBS containing both calcium and magnesium and labeled with 0.5 mg of membrane-impermeable EZ-linkTM Sulfo-NHS-Biotin (Thermo Fisher Scientific, Rockford, IL, USA) per plate for 30 min at room temperature. Plates were washed three times, 5 min each, with 50 mM glycine buffer to quench the reaction. Cells were then washed with PBS lacking calcium and magnesium and lysed with IP Lysis buffer (Thermo Fisher Scientific, Rockford, IL, USA) supplemented with protease inhibitor cocktail kit (Thermo Fisher Scientific, Rockford, IL, USA). Cell lysates were incubated overnight with 90 μL of Dynabeads® MyOne™ Streptavidin C1 (Invitrogen, Carlsbad, CA, USA) on a shaker at 4ºC. The next day beads were collected on a magnet and washed using the following buffers: twice with buffer 1 (2% SDS in dH_2_O), once with buffer 2 (0.1% deoxycholate, 1% Triton X-100, 500 mM NaCl, 1 mM EDTA, 50 mM Hepes pH 7.5), once with buffer 3 (250 mM LiCl, 0.5% NP-40, 0.5% deoxycholate, 1 mM EDTA, 10 mM Tris; pH 8.1), and twice with buffer 4 (50 mM Tris, 50 mM NaCl pH 7.4). To elute proteins and disrupt bead-protein complexes, beads were boiled for 5 min in 60 μL of 1× Laemmli buffer.

### Western Blot

Proteins were first separated with 10% sodium dodecyl sulfate polyacrylamide gel electrophoresis (SDS-PAGE) at 150 V for 1.5 h. The gel was transferred to a nitrocellulose membrane using the Trans-Blot Turbo Transfer System (Bio-Rad Inc., Mississauga, ON, Canada) at 1.3 A and 2.5 V for 7 min. The membrane was washed in PBS buffer and blocked with Intercept® (PBS) Protein-Free Blocking Buffer (LI-COR Biosciences, Lincoln, NE, USA) for 1 h at room temperature. The membrane was then incubated with the primary antibody solution overnight at 4ºC. The following primary antibodies were used for the western blot: rabbit anti-Cx36 (36-4600, Invitrogen, Carlsbad, CA, USA, RRID: AB_2533260) at 1:200, mouse anti-Cx36 at 1:500 (37-4600, Invitrogen, Carlsbad, CA, USA, RRID: AB_253320), mouse anti-β-actin (Sigma-Aldrich Chemie GmbH, Munich, Germany) at 1:1500 concentrations, rabbit anti-phospho-elF2α at 1:1000 (mAB 3398, Cell signaling, RRID: AB_2277684), mouse anti-V5 at 1:1000 (R960-25, Invitrogen, Carlsbad, CA, USA, RRID: AB_2556564) and rabbit tubulin at 1:1000 (MAB12827, Abnova) The secondary antibodies, anti-mouse iRDye 800 and anti-rabbit iRDye 680 (LI-COR Biosciences, Lincoln, NE, USA, RRID: AB_ 271662 & AB_2721181), were used at 1:15,000 concentration and membrane was incubated for 1 h at room temperature. Imaging was performed using the Odyssey® CLx Infrared Imaging System (LI-COR Biosciences, Lincoln, NE, USA). Western blot quantification was performed with ImageJ.

### Dye uptake

HEK293T cells were seeded on 35 mm plates and transfected with Cx36-WT-SNAP or Cx36-C62S-SNAP. SNAP-Cell Oregon Green (New England Biolabs Inc., Boston, MA, USA) was used to label and visualize Cx36 expressing cells. The labeling stock was dissolved in 50 µl of DMSO and diluted to 1:200 in complete clear medium to yield a labeling concentration of 5 µM. Cells were incubated in SNAP-tag labeling medium 37°C, 5% CO_2_ for 30 min. Cells were washed once and incubated in fresh clear medium for 30 min prior to imaging. Dishes were placed in a live-cell imaging chamber at 37°C with CO_2_ and acquired using Zeiss LSM 700 confocal microscope under EC Plan-Neofluar 40x/1.30 Oil M27 objective. Ethidium bromide (EtBr) was added to a final concentration of 10 µM immediately prior to imaging. Images were taken every 15 sec for a duration of 5 min at a 1024 × 1024 pixel resolution. Total EtBr uptake values were used for statistical analysis and were calculated by subtracting the initial fluorescence at T = 0 min from the final fluorescence at T = 10 min. Only cells expressing Cx36-SNAP were used for analysis. Images were analyzed using ImageJ.

### Live/dead assay

HEK293T cells were seeded on 35 mm plates and transfected with Cx36-WT-SNAP or Cx36-C62S-SNAP. ReadyProbes™ Cell Viability Imaging Kit, Blue/Green (Invitrogen, Carlsbad, CA, USA) was used to visualize the total and dead cells 48 h post transfection. Two drops of each stain per 1 ml of media were added to the 35 mm glass-bottom dishes 24 h post transfection. The medium was changed to DMEM lacking phenol red 30 min prior to imaging. Images were taken with a Zeiss LSM 700 confocal microscope using Plan-Apochromat 20x/0.8 M27 objective. Zeiss ZEN imaging software was used to control all imaging parameters.

### Immunocytochemistry

HEK293T cells were briefly washed in PBS and fixed with 2% PFA in PBS for 15 min at room temperature. The coverslips were washed 3 × 10 min with PBS and incubated in the primary antibody (Cx36 1:500, Chemicon MAB3045; p62 1:500, Abcam; Golgi97, 1:100, Thermo Fisher) diluted in 10% normal donkey serum in 0.5% Tx-100 (PBS) overnight at 4°C. At the next day the coverslips were washed with PBS and incubated with the secondary antibodies (From donkey, 1:500, conjugated with Cy3, Alexa 488, Alexa568, or Alexa647; Jackson Immunoresearch, West Grove, PA) diluted in 10% normal donkey serum in 0.5% Tx-100 (PBS). The coverslips were mounted in Vectashield containing DAPI (Vector Laboratories, Burlingame, CA) and sealed with nail polish.

### Imaging

For live cell imaging experiments 2.7×10^5^ HEK293T cells were plated on 30 mm MatTek dishes and 24 h later transfected with 1 µg of Cx36-SNAP. At the next day, these cells were treated with 3 µM SNAP-Cell TMR-Star (NEB) for 30 min at 37°C and washed in 10% FBS DMEM for 30 min. Afterwards the DMEM was substituted with Ames medium (Sigma). Live cell imaging experiments were conducted with a Nikon Eclipse Ti microscope using a 60x objective. After TMR labeling HEK293T cells were placed in an incubation chamber kept at 37°C, 5% CO_2_ and imaged for 30 min in 30 sec intervals. Fluorescence images were acquired with a Zeiss LSM800 confocal microscope using an 60x objective with a numerical aperture of 1.3.

### Image analysis

The frequency and size of connexin whorls were measured with ImageJ. The number of whorls was quantified for individual cells (z-stack of 10 × 0.2 µm) using the cell counter plugin. Intracellular vesicles without a recognizable lumen were excluded from quantification. These were usually vesicles with an inner diameter smaller than 0.3 µm. A linear ROI and the measure function were used to determine the inner diameter of individual whorls. For colocalization analysis of Cx36 and Golgi97, cells were scanned with a Leica TCS SP8 confocal microscope using a 63x/1.4 oil objective, at a pixel size of ∼45 nm. Image stacks (∼ 8 µm) were subsequently processed in Fiji (Schindelin et al., 2012): First, stacks were thresholded using the *Moments* algorithm of the *Auto Threshold* function. Then, colocalized pixels were identified using the *Colocalization Highlighter*. In the resulting 8-bit image, four regions of interest (ROI = 3.01 × 3.01 μm^2^) were placed on the colocalized puncta and the colocalized area was measured using the *Analyze Particle* function which was set to count only puncta larger than 0.125 µm^2^. Subsequently, the same ROIs were placed on the Golgi97 image stacks and again, the stained area was measured. From these data, the percentage of colocalization with Cx36 was calculated and averaged for each transfected construct (four ROIs analyzed in each stack, three stacks per coverslip, two coverslips per construct). In total, three independent transfection experiments were analyzed and the resulting data were compared by a two tailed Mann-Whitney test.

### Conventional correlative light and electron microscopy (CLEM)

HEK293T cells (1×10^5^) were seeded in 35 mm µ-Dishes with a gridded polymer coverslip bottom (Cat.No. 81166; Ibidi, Gräfelfing, Germany) coated with 0.01% poly-L-lysine. Next day cells were transfected (FuGene) with either pCx36-SNAP or pCx36 C62S-SNAP. On the third day cells were stained with 100 nM SNAP-Cell TMR-Star (New England BioLabs) for 30 min at 37°C and chemically fixed with a mixture of 4% paraformaldehyde and 0.1% glutaraldehyde in 0.2 M HEPES buffer, pH 7.4, for 15 min at 37°C. ROIs were observed with a confocal laser scanning microscope (Leica TCS SP5) and the software LAS AF (Leica). After imaging cells were fixed with 2.5% glutaraldehyde (Science Services) in 0.2 M HEPES, pH 7.4, for 1 h at RT and postfixed with 1% osmium tetroxide (Science Services) in 0.1 M cacodylate buffer, pH 7.4, for 2 h on ice. After washing, cells were dehydrated in a graded ethanol series (30%, 50%, 70%, 90%) and finally two rinses in anhydrous ethanol and three rinses in anhydrous acetone at RT. Cells were infiltrated and flat-embedded in graded mixes of acetone and Epon 812 (Sigma). After resin polymerization the gridded polymer bottom was removed, and the coordinates were transferred to the resin surface allowing trimming. Serial 70 nm sections were cut with an ultramicrotome (Leica EM UC7RT) and collected on formvar-coated EM copper slot grids. After automated staining with uranyl acetate and lead citrate (Leica EM AC20), samples were observed via either Zeiss TEM 902 A (50 kV, TRS slow-scan 2K CCD camera, software ImageSP by TRS image SysProg) or Jeol TEM JEM2100plus (200 kV, emsis XAROSA CMOS TEM camera, software RADIUS by Emsis). Stitching and overlay of CLSM and TEM images was done using Photoshop (Adobe).

### Serial Block-Face Scanning Electron Microscopy (SBF-SEM) CLEM

After light microscopy, HEK293T cells selected for SBF-SEM were fixed in 2.5% glutaraldehyde in 0.1 M cacodylate buffer, pH 7.4. Subsequently, samples were processed via adapted version of the NCMIR rOTO-post-fixation protocol (Deerinck et al., 2010) and embedded in hard Epon resin, ensuring pronounced contrast and electron dose resistance for consecutive imaging. All procedures were performed on the Ibidi µ-Dish. In brief, after fixation, samples were post-fixed in 2% osmium tetroxide (Science Services) and 1.5% (w/v) potassium ferrocyanide (Riedel de Haën) in cacodylate buffer for 1 h on ice. The cells were then incubated in 1% (w/v) thiocarbohydrazide (Riedel de Haën) in dH_2_0 for 20 min, followed by an additional 2% osmication step in water for 30 min at room temperature. After washing in dH_2_0, samples were submerged in 1% aqueous uranyl acetate overnight at 4°C. Cells were then incubated in freshly prepared Walton`s lead aspartate (Pb(NO_3_)_2_ (Carl-Roth), L-Aspartate (Serva), KOH (Merck)) for 30 min at 60°C. Subsequently, cells were dehydrated through a graded ethanol (Carl-Roth) series (30%, 50%, 70% and 90%) on ice for 7 min each, before rinsing in anhydrous ethanol twice for 7 min and twice in anhydrous acetone (Carl-Roth) for 10 min at room temperature. Afterwards, cells were infiltrated in an ascending Epon:acetone mixture (1:3, 1:1, 3:1) for 2 h each, before an additional incubation in hard mixture of 100% Epon 812 (Sigma). Final curation was carried out in hard Epon with 3% (w/w) Ketjen Black (TAAB) at 60°C for 48 h. Once polymerized, the µ-Dish bottom was removed via toluene melting from the resin block including the attached cells and finder grid imprint. ROIs were trimmed and the sample blocks were glued to aluminium rivets using two-component conductive silver epoxy adhesive and additionally coated in a 30 nm thick gold layer. The rivet containing the mounted resin block was then inserted into the 3View Gatan stage, fitted in a Jeol JSM 7200F, and aligned parallel to the diamond knife-edge. The cells proved to be stable under imaging conditions of 3.1 kV accelerating voltage, high vacuum mode of 10 Pa, utilizing a 30 nm condenser aperture and a positive stage bias of 600 V. Imaging parameters were set to 6 nm pixel size, 3.4 µs dwell time, in between ablation of 60 nm and an image size of 10240×10240 pixels. Overall, an approximate volume of 60×60×19 µm (à 320 slices) was acquired. Image acquisition was controlled via Gatan Digital Micrograph software (Version 3.32.2403.0). Further post-processing, including alignment, filtering and segmentations were performed in Microscopy Image Browser (Version 2.7,(Belevich et al., 2016)). To keep the computational resources to a manageable limit, the final stacks were binned before exporting the files for volumetric visualization in Amira 3D (Version 2021.1, Thermo Fisher).

### Statistical analysis

Statistical analysis and data presentation were performed using GraphPad Prism 8. Values reported consist of mean ± SEM. Normality was tested using the Anderson-Darling and the D`Agostino-Pearson test. Significance was tested using a two-tailed t-test, two-tailed Mann-Whitney U test or a Kruskal-Wallis test for multiple comparisons.

## Results

### Removal of the PDZ binding motif of Cx36 causes ER retention

Previous studies have shown that the PDZ binding motif in Cx36 is essential for the formation of electrical synapses. Fusion proteins such as Cx36-EGFP, in which an EGFP tag is fused to the C-terminal tip of Cx36, exhibit severe transport deficits and tend to accumulate in the cytoplasm (Helbig et al., 2010). C-terminal tags are likely to compromise the functionality of the PDZ binding motif because they block access to the extreme C-terminus. Currently, it is unknown how exactly the inhibition of this motif affects the intracellular transport of Cx36 and why it results in the intracellular accumulation of the connexin. To shed light on the processes causing these transport deficits, we generated a Cx36 mutant that lacks the PDZ binding motif and a Cx36-SNAP construct in which a SNAP-tag is positioned at the C-terminus (Figure 1A). In an initial experiment we expressed these constructs in HEK293T cells and compared their distribution to wild type Cx36. The deletion of the PDZ binding motif had a striking effect on the distribution of the Cx36/S318ter mutant. This mutant did not assemble into compact perinuclear structures but instead formed large intracellular vesicles that resembled annular junctions (Figure 1B). A similar phenotype was observed for the Cx36-SNAP construct. We quantified the frequency and size of these vesicles (here and throughout the paper referred to as whorls) and found that both parameters were significantly increased in the mutant and the fusion protein (Figure 1C-D). None of the mutants influenced the frequency of gap junction clusters, whereas deletion of the PDZ binding motif had a striking effect on the intracellular distribution of the Cx36/S318ter mutant and the Cx36 SNAP construct (Figure 1E). The Cx36/S318ter displayed around 1.3 whorls per cell and the Cx36-SNAP construct had even more than twice as many. The size of these whorls was also increased in comparison to the wild type but there were no size differences between Cx36/S318ter and Cx36-SNAP (Figure 1D). During image acquisition, we observed that Cx36 whorls were often quite bright and displayed intensities in the same order of magnitude as gap junctions, which suggests that they contain a high concentration of connexin molecules. One possibility to explain the whorl formation that is induced by the Cx36 mutants is an increased internalization rate. However, Cx36 whorls were rarely associated with gap junctions, but they instead localized around nucleus and in the cytoplasm. This suggests that these structures are formed as a result of a premature release mechanism but not as an internalization event. To identify the subcellular compartment from which these vesicles originate, we analyzed the colocalization of Cx36, Cx36/S318ter and Cx36-SNAP with ER and Golgi markers. Here we observed a striking colocalization of Cx36 whorls and the ER membrane marker GFP-Sec61B (Figure 1F). On the contrary we didn`t find any obvious differences in the association with the Golgi apparatus for Cx36 and the Cx36/S318ter mutant. These constructs rarely colocalized with the trans Golgi marker Golgin-97 (Figure 1G). The lack of association with the Golgi and the extensive colocalization of Cx36 whorls with GFP-Sec61B suggest that the disruption of the PDZ binding motif prevents Cx36 from exiting the ER. The resulting formation of connexin whorls is likely to reflect a spillover mechanism that induces a premature release of connexins from the ER.

**Figure 1.**
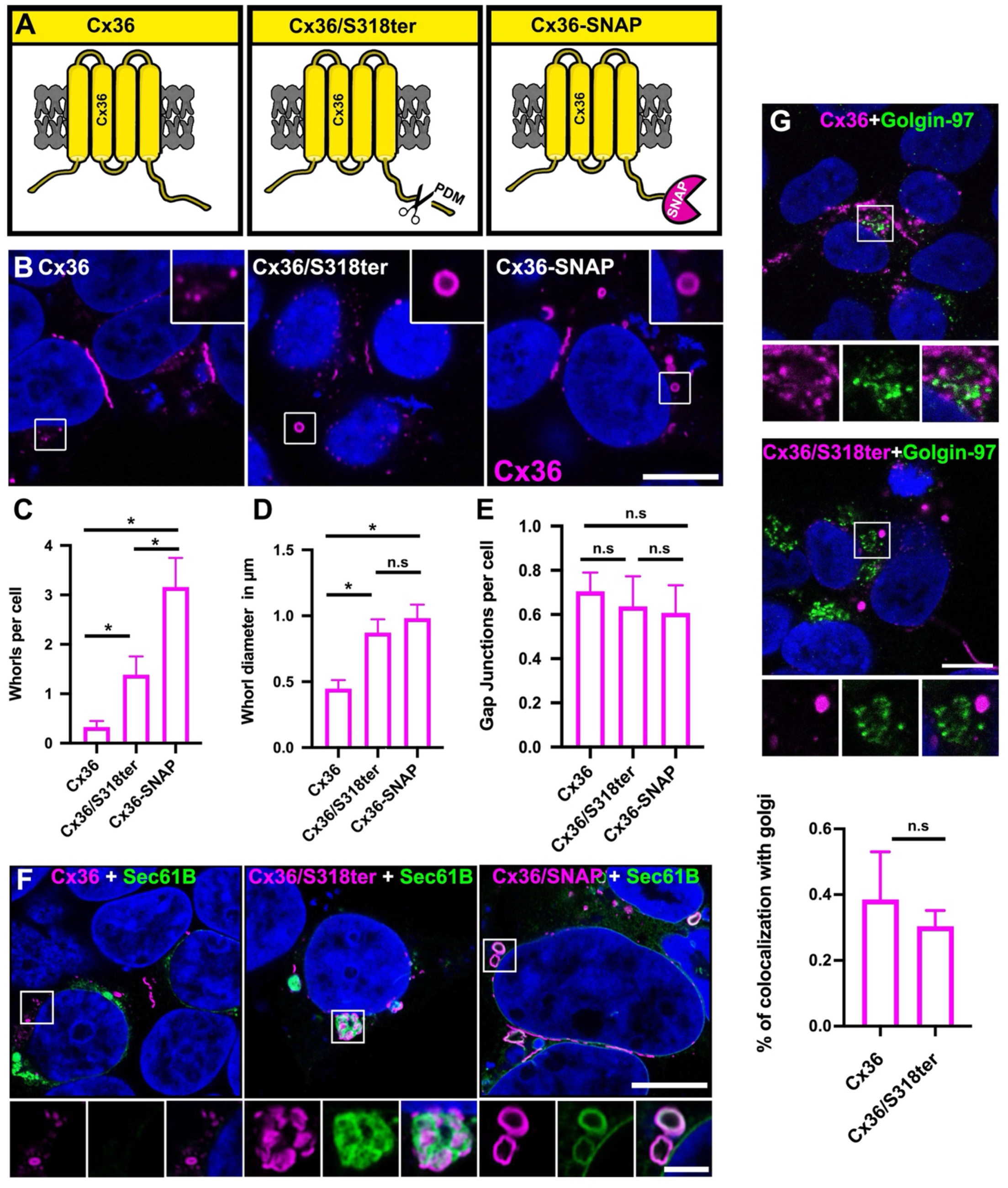
Removal of the PDZ binding motif of Cx36 and C-terminal SNAP-tag leads to the formation connexin whorls. **(A)** Illustration of different Cx36 mutants used in this study. **(B)** Perinuclear distribution of wild type Cx36 in transfected HEK293T cells. Removal of the PDZ binding motif or the addition of a C-terminal SNAP in Cx36 results in the formation of large annular inclusions (whorls). Scale: 10 µm. **(C)** Frequency of intracellular whorls per cell. Whorls were counted in 9-11 cell clusters from two independent transfections. Compared to WT Cx36, Cx36/S318ter and Cx36-SNAP transfected cells show a significant increase in whorl formation. P<0.05, *. Values are Mean with SEM and significance was determined by a Kruskal-Wallis test followed by a post-hoc Dunn`s analysis. **(D)** Average size of connexin whorls in all three Cx36 variants. Cx36 whorls are increased in size in Cx36/S318ter and Cx36-SNAP transfected cells. 11-30 whorls were measured per condition. P<0.05 *. Significance was tested using a Kruskal-Wallis followed by a post-hoc Dunn`s analysis. **(E)** Frequency of gap junctions per cell. P>0.05, N.S not significant. 10-11 cell clusters from two independent transfections were quantified per condition. Values are Mean with SEM and significance was determined by a Kruskal-Wallis test followed by a post-hoc Dunn`s analysis. **(F)** Colocalization of Cx36 whorls and the ER membrane marker Sec61B-GFP. Scale: 10 µm. **(G)** Cx36 and Cx36/S318ter show no differences in association with the Golgi marker Golgin-97. P=0.1345, N.S not significant. Scale: 10 µm. Colocalization was quantified in an entire stack. Values are mean ± SEM and were obtained from three independent transfection experiments. Significance was determined by a two tailed Mann-Whitney test.

### Extracellular cysteines are required for whorl formation

Previous studies have shown that overexpression of GFP-tagged ER resident proteins results in the generation of multilamellar vesicles that consist of stacked ER membranes. The formation of these structures is triggered by dimerization of GFP. Homophilic interactions of multiple GFP monomers along ER tubules cause the ER to coil up into concentric whorls, eventually leading to their release (Snapp et al., 2003). As connexins have the ability to self-assemble into dense aggregates, we reasoned that a similar mechanism is responsible for the whorl formation we observed here. A premature docking of opposing connexons in the ER might have the same effects and result in a structural reorganization of ER membranes. This mechanism could explain the increased whorl formation we observed for ER retained Cx36 mutants. To test if the formation of connexin whorls requires a gap junction-like docking mechanism, we substituted individual cysteines (Cysteine 55 and 62, Figure 2A) in the extracellular loop of Cx36 with serines. Previous studies have shown that mutations in this region prevent gap junction formation because they interfere with the intercellular docking of connexons (Placantonakis et al., 2002). Eliminating the extracellular cysteines should also prevent the formation of ER whorls, provided a similar docking mechanism is responsible. To test this hypothesis, we expressed Cx36 cysteine mutants in HEK293T cells and quantified the frequency of whorls for each construct. As expected, the expression of these mutants entirely prevented gap junction formation but also resulted in a striking reduction of ER whorls (Figure 2C-D). For each of the cysteine mutants, we found hardly any whorls. This difference was most distinctive for the ER retained Cx36/318ter and the Cx36-SNAP construct (Figure 2E). These constructs displayed the highest number of whorls. Each of the corresponding cysteine mutants, however, formed hardly any whorls, indicating that the formation of connexin whorls requires intact extracellular loops. Surprisingly we found that all cysteine mutants showed a marked decrease in protein expression (Figure 2B) and apparent signs of cell damage. Especially, cells that expressed larger amount of the mutant often had a rounded shape and deformed nuclei.

**Figure 2.**
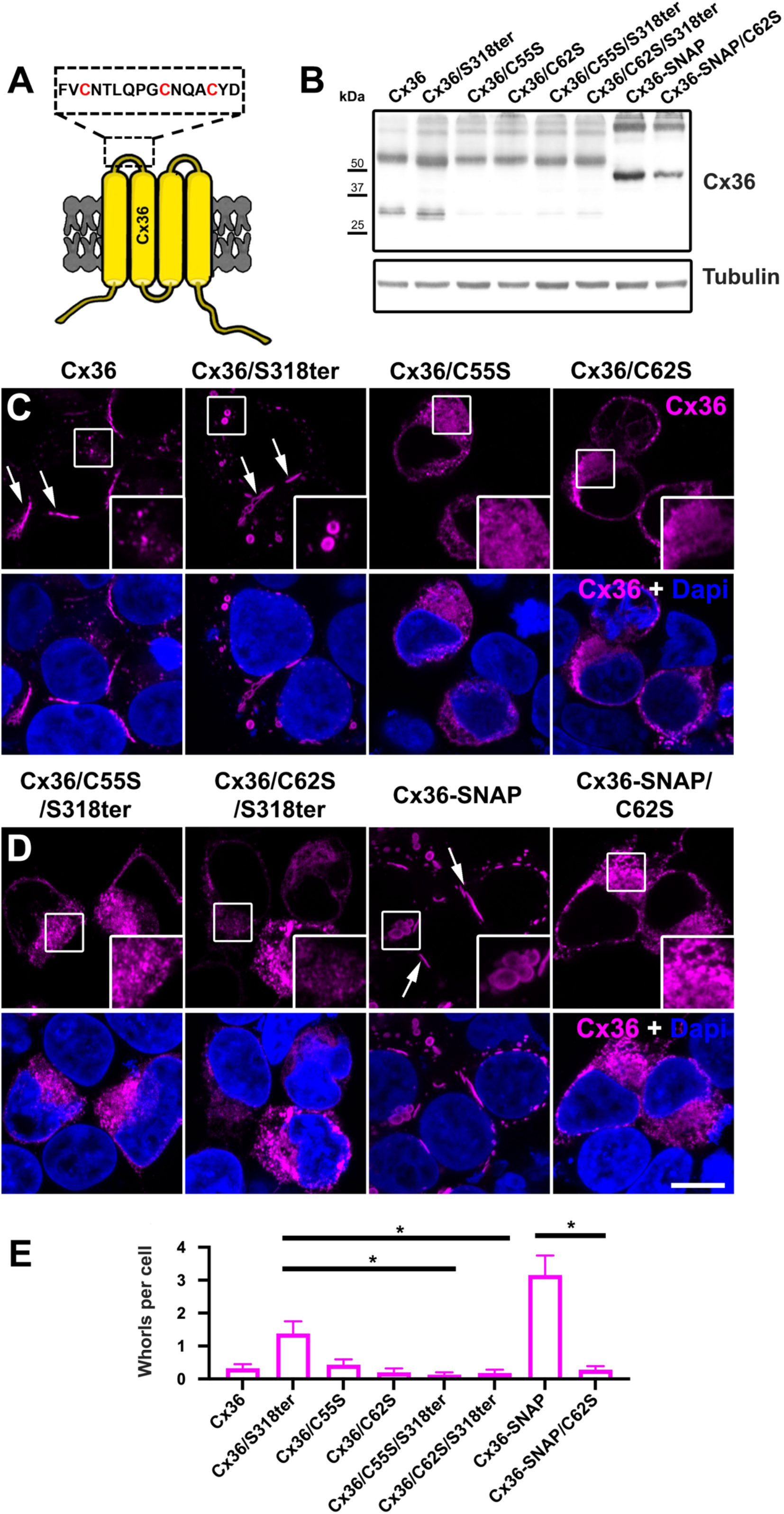
Mutations of cysteines in the extracellular loop of Cx36 prevent whorl formation. **(A)** Cartoon illustrating the positions of the extracellular cysteines in Cx36. **(B)** Protein levels of cysteine mutants in transfected HEK293T cells. Each of the cysteine mutants exhibits a marked reduction in Cx36 expression. **(C)** HEK293T cells expressing wild type Cx36 and the Cx36/S318ter mutant. Substitution of cysteines 55 or 62 result in a diffuse distribution of Cx36 in the cytoplasm. **(D)** Cysteine mutations in the Cx36/S318ter or the Cx36-SNAP construct prevent whorl formation. Images are presented as maximum projections of 10 slides (2 µm). **(E)** Frequency of intracellular whorls per cell. Whorls were counted in 9-11 cell clusters from two independent transfections. Compared Cx36/S318ter and Cx36-SNAP the corresponding cysteine mutants show a significant decrease in whorl formation. P<0.05. Values are Mean with SEM and significance was determined by Kruskal-Wallis test followed by a post-hoc Dunn`s analysis. Scale: 10 µm.

### Ultrastructure of ER whorls

Our initial experiments have shown that the formation of connexin whorls requires the extracellular loop cysteines. Provided that Cx36 whorls are formed by a gap junction-like docking mechanism, it is to be expected that they are identical to actual gap junctions in terms of structure and density. To test this hypothesis, we resolved the ultrastructure of ER whorls and gap junctions in Cx36-SNAP transfected HEK293T cells using a correlative light and electron microscopy approach. Cx36-SNAP expressing cells were labeled with TMR and imaged in a live cell imaging set up. Individual regions of interest (ROI) were then traced back and scanned with the high resolution of EM, which allowed us to compare the ultrastructure of whorls and gap junctions (Figure 3A_i-iii_). As expected, we found that Cx36 whorls consisted of multiple membranes (Figure 3B_i_). Each sheet within these whorls contained two parallel membrane layers with densely packed connexins. This configuration had obvious features of gap junctions, which suggests that ER whorls are indeed formed by a similar head-to-head docking mechanism. Interestingly, we also observed that some gap junctions were associated with stacks of intracellular membranes that seemed to consist of docked channels (Figure 3B_ii_). The configuration of this structure was indistinguishable from the gap junction.

**Figure 3.**
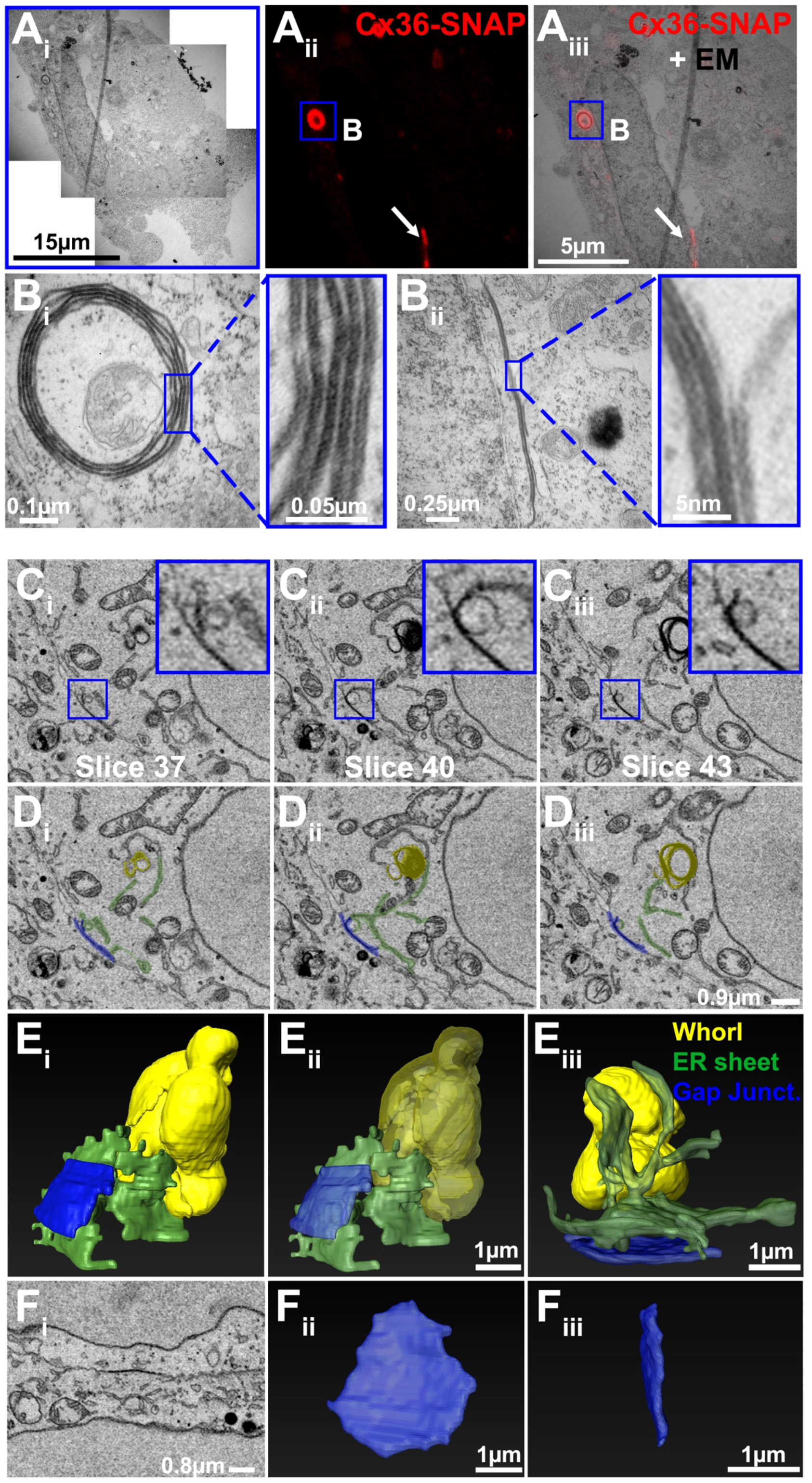
CLEM and SBF-SEM reveal the ultrastructure of connexin whorls. (A_**i**_**-B**_**iii**_) Fluorescence signals were correlated with electron micrographs. These experiments revealed that connexin whorls are multimembrane vesicles. Each layer within these vesicles consisted of two adjoined membranes. This configuration was indistinguishable from the structure of gap junctions. **(C**_**i**_**-D**_**iii**_) Single slices showing electron micrographs of gap junction (blue), ER whorls (yellow) and ER sheets (green). ER whorls are directly connected to gap junction via membranes of the ER. **(E**_**i**_**-F**_**iii**_) 3D reconstruction of gap junctions and ER whorls via serial block face imaging. Note that ER sheets are continuous with ER whorls and come in close contact with the gap junction. Length of scale bar is indicated in each panel.

To further characterize the structure of Cx36 whorls we reconstructed the entire volume of individual vesicles using serial block-face SEM imaging. These reconstructions illustrated the ellipsoid shape of Cx36 whorls (Figure 3E_i_ & 3E_iii_) and their complex multi membrane structure (Transparent version in figure 3E_ii_). In line with the CLEM data, we observed direct conjunctions between ER sheets, gap junctions and ER whorls. We labeled these structures with pseudo colors in single SBF-SEM sections (Figure 3C_i_-D_iii_, blue: gap junction, yellow: whorl, green: ER sheet). These scans showed that ER sheets were often aligned with the gap junction and simultaneously contacting the whorl. Interestingly, we observed a single vesicle that was seemingly about to be removed from the gap junction forming a continuous connection with an ER sheet (Figure 3C_ii_-D_ii_; insets in C_ii_).

### Pulse-chase labeling experiments reveal the mechanism of whorl formation

So far, our data have shown that the ability of connexins to self-assemble into dense aggregates serves as a basis for a premature ER release mechanism. The proposed docking mechanism, however, does not explain why connexin aggregates assemble into compact vesicles. To understand the structural changes underlying whorl formation we conducted live cell imaging experiments. Cx36-SNAP transfected cells were treated with the SNAP specific TMR ligand and monitored in a live cell imaging set up. During an acquisition time of 30 min, we did not detect any obvious structural changes of individual whorls in the region of interest, except for minor movements (Figure 4A). Interestingly, we recorded a vesicle that appeared to be removed from the gap junction (Figure 4A, magnified inset). This structure was not internalized from the center of the gap junction like an annular junction, but it was seemingly coiling up along the cell membrane. Considering our previous observation, it seems possible that this structure represents a whorl that is about to be formed at the gap junction. To further resolve the time course of whorl formation with improved spatial resolution we performed pulse chase labeling experiments using two different SNAP ligands. The SNAP-tag can be conjugated with interchangeable fluorescent dyes that form irreversible bonds with the enzyme (Cole, 2013). This technology is well suited for pulse-chase experiments because it allows a differential labeling of Cx36-SNAP proteins that were synthesized at different time points. As a pulse we incubated Cx36-SNAP transfected cells with the SNAP-Cell Oregon Green ligand. This initial incubation labels all preexisting SNAP molecules prior to the chase. At different time points after the initial incubation, we applied the TMR ligand to label Cx36-SNAP molecules that where synthesized after the pulse. This strategy allowed us to distinguish two different generations of Cx36-SNAP proteins using confocal scans. The pulse-chase labeling experiment revealed that ER whorls consist of old and newly synthesized Cx36 proteins (Figure 4B). We often observed that new material was incorporated to the lumen of preexisting whorls. In line with the SBF-SEM data, we found newer TMR labeled vesicles in direct association with older Oregon Green labeled gap junctions. We scanned a single whorl after a 30 min TMR chase using the high resolution of the Airy scan. Here, we found that Cx36 whorls consisted of several layers (Figure 4C), which is consistent with our previous observation. Newly synthesized material (TMR labeled) always covered the inside of the whorl whereas older, Oregon Green labeled material formed the outer layer. These observations suggest that connexin whorls are not isolated vesicles, as for instance connexosomes (Falk et al., 2016), but they appear to be dynamic structures that are constantly growing. Given the direct connections between gap junctions and ER sheets, it is likely that a certain percentage of Cx36 whorls is immediately formed at the cell membrane, creating the false impression of an internalization event.

**Figure 4.**
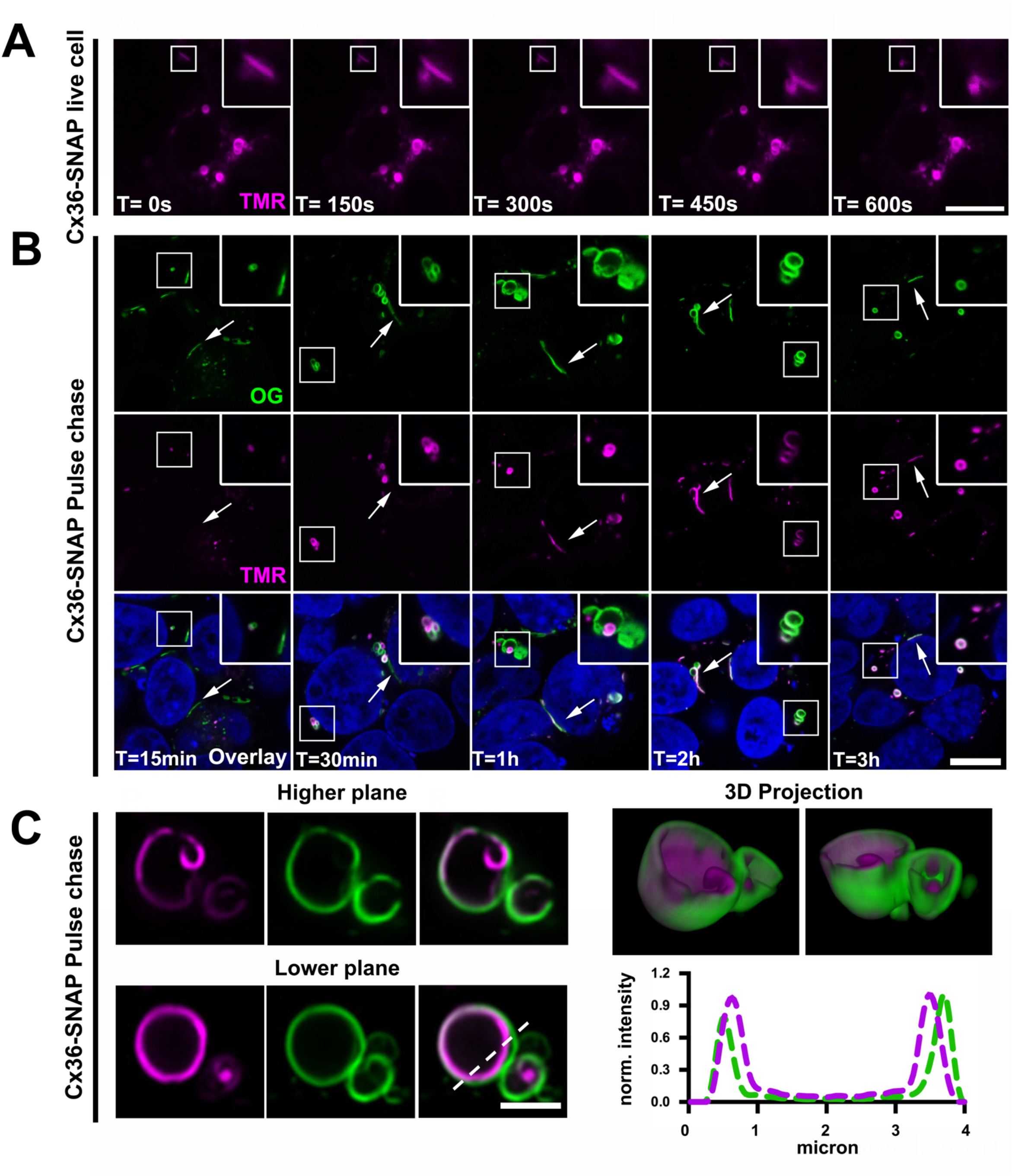
Live cell imaging and pulse chase experiments reveal the mechanism of Cx36 whorl formation. **(A)** Cx36-SNAP expressing HEK293T cells were labeled with TMR and imaged in a live cell imaging chamber. Images are shown in 5 min intervals. Scale: 5 µm. **(B)** Cx36-SNAP transfected HEK293T cells were pulsed with Oregon Green and chased with TMR at different time points: 15 min, 30 min, 1 h, 2 h, 3 h. Scale: 10 µm. **(C)** Airy scan of pulse-chased whorl. Two different planes are shown. TMR labeled material (magenta) is covering the inside of the whorl. A 3D reconstruction demonstrates that older Oregon Green labeled protein is forming the outer layers of the whorl. Scale: 2.5 µm.

### Characterization of cysteine mutants

Our experiments suggest that overexpression of Cx36 cysteine mutant forms results in apparent cell damage causing morphological changes. This raises the question of whether disrupting whorl formation is toxic for transfected HEK293T cells. As connexin whorls function as a removal mechanism, we reasoned that this cell damage is caused by an overabundance of the cysteine mutants accumulating in the ER. To test this, we co-transfected the ER retained Cx36/S318-ter, Cx36-SNAP and the corresponding cysteine mutants with the luminal ER marker DsRed2 ER5. We found that each of the cysteine mutants displayed strong colocalization with DsRed2 ER5 (Figure 5A_ii_, 5A_iv_ 5B_ii_, 5B_iv_). Due to the low expression of these mutants, we had to increase the laser intensity in these conditions. A similar degree of colocalization was observed when Cx36/S318-ter and Cx36-SNAP transfected cells were imaged with the same laser intensities. However, we worked with lower intensities in these conditions to avoid an oversaturation of connexin whorls. Although Cx36 whorls originate from the ER, they didn`t seem to colocalize with the luminal ER marker DsRed2 ER5 (Figure 5A_i_, 5A_iii_, B_i_, B_iii_). This can be explained by the fact that docking of connexins in the ER lumen displaces the luminal content leaving no space for the ER marker protein (Lichtenstein et al., 2009). To determine the extent of cell death we carried out cell vitality assays. A live/dead assay was used to visualize dead cells. Although, Cx36/C62S-SNAP transfectants often appeared to be damaged based on cell morphology and shape of their nuclei, our live/dead assay showed no apparent differences in the fraction of dead cells for each condition (Figure 5C_i_). In Cx36-SNAP and Cx36/C62S-SNAP we observed around 10% of dead cells. This number was not affected by the cysteine mutant. We also tested for upregulation of ER stress markers and determined the level of phosphorylated eukorayoric initiation factor alpha (elf2α) in cells that were transfected with the cysteine mutant (Figure 5C_ii_). Here we didn`t observe any changes in the phosphorylation state of elf2α when we compared Cx36-SNAP and Cx36/C62-SNAP expressing cells. As a positive control we induced ER stress applying 2 mM DTT for 1h. This treatment, unlike overexpression of Cx36/C62-SNAP, caused an increase in phosphorylation of elf2α.

**Figure 5.**
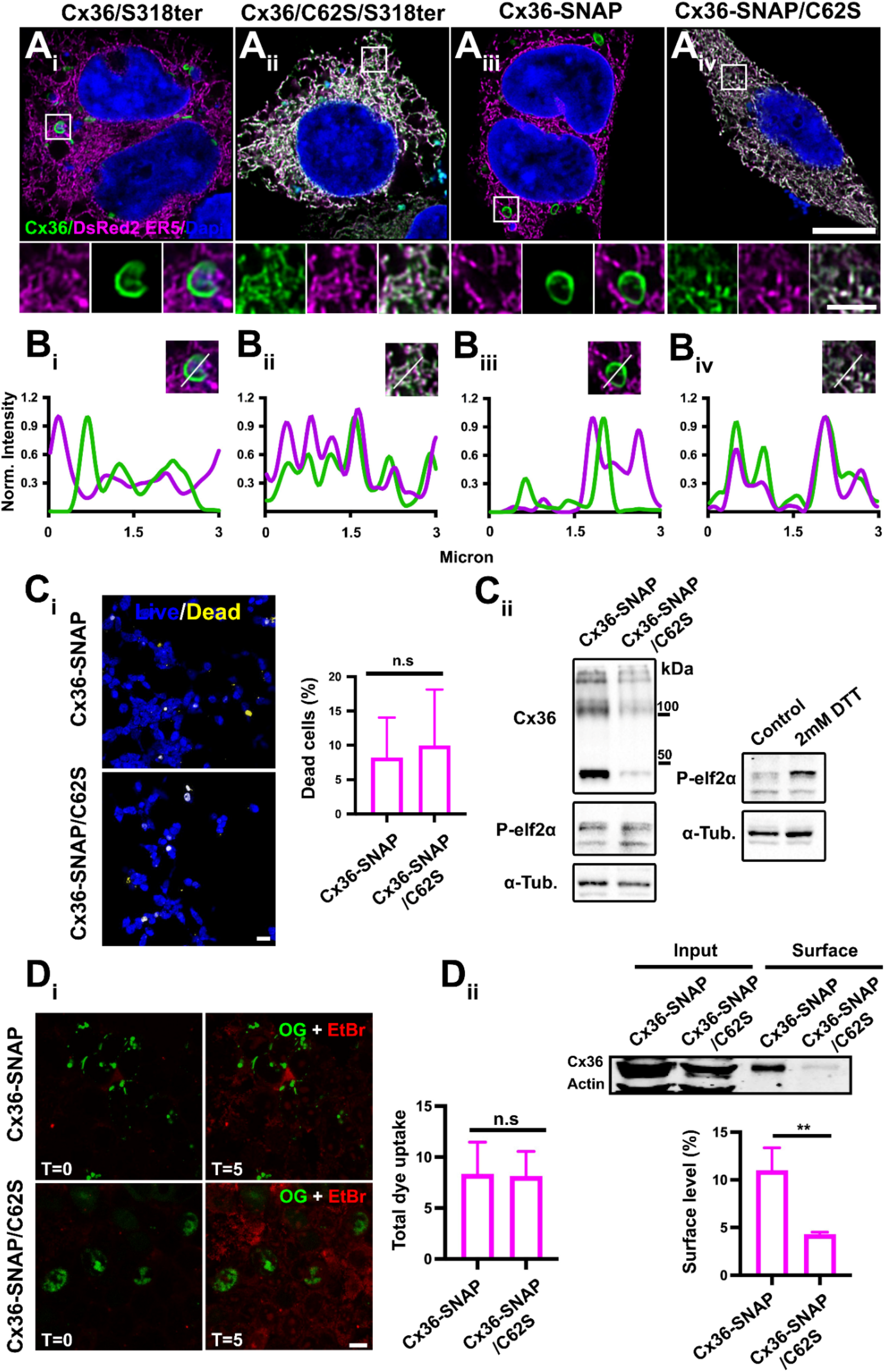
Characterization of Cx36 cysteine mutants. **(A**_**i**_**-A**_**iv**_) Cotransfections of Cx36 mutants and ER localized DsRed2ER5 reveal that the cysteine mutants Cx36/C62S/S318ter and Cx36-SNAP/C62S are almost entirely confined to the ER. Whorls formed by Cx36/S318ter or Cx36SNAP transfectants do not colocalize with dsRed2ER5. Scale: 10 µm. Magnified insets: 2 µm. **(B**_**i**_**-B**_**iv**_) Line scan depicting the intensity along each ROI. Intensity profile of DsRed2ER5 (magenta) and the cysteine mutants (green) overlaps entirely. **(C**_**i**_) Representative confocal images of Cx36-WT-SNAP and Cx36-SNAP/C62S transfected cells stained with live/dead viability staining kit. NucBlue® Live reagent: stains the nuclei of all cells; detected with a standard DAPI filter. NucGreen® Dead reagent: stains only the nuclei of dead cells; detected with standard GFP (green) filter set. Scale: 20 µm. Quantification of dead cells 24 h post staining with live/dead dyes. Plot with SEM; Mann-Whitney test (two-tailed), N.S. not significant. **(C**_**ii**_) Lysates of Cx36-SNAP and Cx36-SNAP/C62S transfected HEK293T cells were tested for phosphorylated elf2α which serves as an ER stress marker. No changes in elf2α phosphorylation were detected between Cx36-SNAP and Cx36-SNAP/C62S transfected cells. As a positive control we induced ER stress in non-transfected cells using 2 mM DTT for 60 min. This treatment resulted in increased phosphorylation of elf2α. **(D**_**i**_) EtBr dye uptake assays demonstrate that substitution C62S doesn’
st affect the permeability of transfected HEK293T cells. Representative confocal images of EtBr uptake in Cx36-SNAP and Cx36/C62S-SNAP expressing cells labeled with SNAP-Cell Oregon Green dye. Scale bar: 10 µm. Bar graph quantification of total increase in fluorescence 5 min after EtBr application. Only cells expressing Cx36 were used for dye uptake analysis. Plot with SEM; Mann-Whitney test (two-tailed), N.S. not significant. **(D**_**ii**_) Cell surface biotinylation assay displaying levels of Cx36-SNAP/C62S and Cx36-SNAP at the membrane. Total cell lysates (input) show expression of Cx36 and intracellular ß-actin. The streptavidin pull-down fractions (surface) show a reduction in membrane levels of Cx36-SNAP/C62S when compared to Cx36-WT. ß-actin served as an internal control. Bar graph with mean and SEM; unpaired t test (two tailed), **< 0.01. n=3 experiments.

Another possibility to explain the increased cell damage we have observed for the cysteine mutants is the formation of leaky hemichannels. Previous studies have shown that disruption of extracellular cysteines in Cx43 is correlated with an increase in hemichannel activity (Retamal et al., 2007). It is possible that the C62S substitution has similar effects on the permeability of Cx36 connexons and leads to aberrant channel activity resulting in increased uptake of extracellular Ca^2+^, which could be sufficient to induce apoptosis. To exclude this possibility, we measured the channel activity of the cysteine mutant and compared the ethidium bromide uptake of Cx36-SNAP and Cx36/C62S-SNAP transfectants. To identify cells that express the fusion proteins, we incubated the samples with SNAP Cell Oregon Green prior to the measurement. The ethidium bromide assay showed no differences in dye uptake between the two conditions (Figure 5D_ii_). For the two constructs we observed the same of extent dye uptake after 5 min of perfusion, which is likely to originate from endogenous hemichannel activity. In line with these observations, surface biotinylation assays showed a relative decrease in the surface expression of the Cx36/C62S-SNAP mutant (Actin was used as a negative control to exclude cytoplasmic contaminants) (Figure 5D_ii_). This confirms that aberrant hemichannel activity of plasma membrane localized Cx36 mutants is unlikely to induce the cell damage we have seen in prior experiments. Although the overall dye uptake was similar for both constructs, we cannot exclude that the C62S mutation influences the permeability of Cx36 connexons since the Cx36/C62-SNAP mutant exhibited reduced surface expression, making direct comparisons impossible.

### Cx36 whorls associate with the p62 protein

The preceding experiments have drawn a precise image of how Cx36 is assembled into ER whorls. To understand how these structures are further processed and potentially degraded we carried out a BioID screen with a recently described dGBP-TurboID construct, consisting of a conditionally destabilized GFP nanobody (dGBP) and the TurboID ligase. This strategy can be used to target the biotin ligase to GFP labeled proteins of interest (Xiong et al., 2021). In order to specifically target whorl incorporated Cx36 channels, we coexpressed Cx36-EGFP and Cx36-SNAP. This strategy allowed us to direct the TurboID enzyme to large connexin whorls that were formed by the two fusion proteins (Figure 6A). To induce proximity biotinylation we treated transfected cells with 50 µM Biotin for 3 h and performed the pull-down afterwards. The pull-down results were confirmed by testing streptavidin reactivity in the eluted samples (Figure 6A). A sister gel was incubated with a V5 antibody (V5 epitope tag on the N-terminus of dGBP-TurboID) to visualize the isolated fusion protein. A Cx36 antibody was used to confirm streptavidin affinity capture of Cx36-SNAP and Cx36-EGFP. Both proteins were highly abundant in the precipitates of the pulldown and occurred as a double band (Upper band: Cx36-EGFP, lower band: Cx36-SNAP). To further confirm the correct targeting of Cx36 whorls, we tested the biotin labeling in co-transfected cells. Here, we observed that V5-dGBP-TurboID and Cy3-Streptavidin labeling (Figure 6B.) was entirely confined to Cx36-EGFP containing ER whorls. The eluates of the BioID assay were subjected to a mass spectrometry to screen for potential interactors of Cx36. Among the proteins that were identified in the precipitates (Figure 6C), we detected the p62 protein (the p62 protein was also present in the negative control, but as it has a ubiquitous function for protein degradation, it is not surprising, that it occurred in a complex with V5-dGBP-TurboID). This finding was of particular interest for us as p62 is a known player of ER phagy, a type of autophagy that degrades fragments of the ER to remove potential stressors. To validate these results and to confirm an association between Cx36 and p62, we analyzed the localization of these proteins in Cx36-SNAP transfectants. In this experiment, we observed extensive colocalization between p62 and Cx36-SNAP (Figure 6B). P62 immunoreactivity was closely aligned with the ER whorl. P62 functions as an adapter protein that targets ubiquitinated protein into autophagosomes (Pankiv et al., 2007). This suggest that ER whorls represent a point of no return from which Cx36 whorls are subjected to the proteolytic systems of the cell.

**Figure 6.**
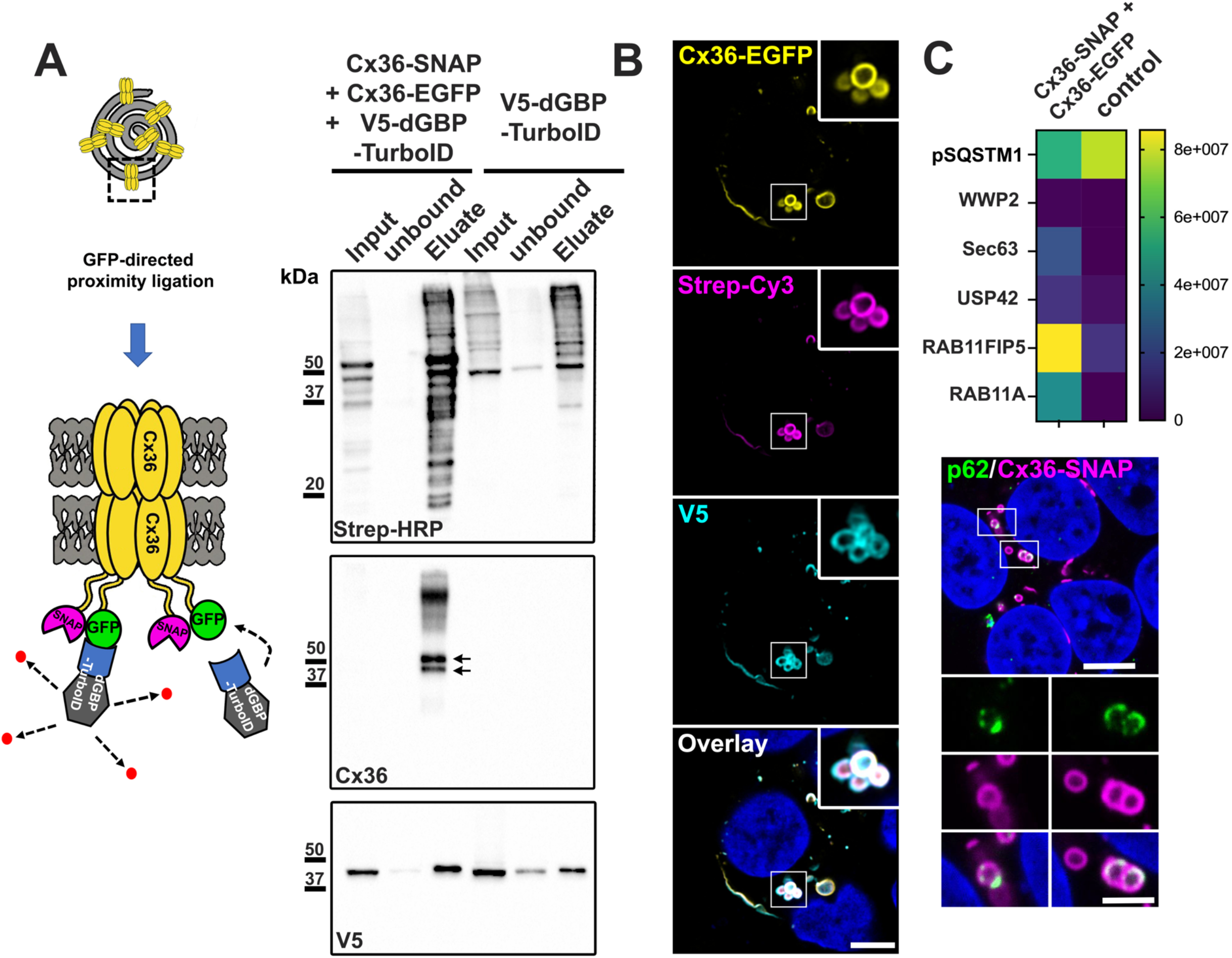
Cx36 associates with the p62 protein. **(A)** Cartoon illustrating whorl specific proximity ligation. Co-expression of Cx36-SNAP and Cx36-GFP leads to the incorporation of Cx36-GFP into large connexin whorls. The GFP-tag on Cx36-EGFP functions as a bait to direct the V5-dGBP-TurboID to connexin whorls, allowing biotinylation of whorl associated proteins. Western blots confirming streptavidin affinity capture. Streptavidin-HRP was used to detect biotinylated proteins. A V5 antibody was used to detect the V5-dGBP-TurboID construct. Cx36-EGFP and Cx36-SNAP were co-isolated. The Cx36 antibody detects two bands in the eluates of Cx36-EGFP+ Cx36-SNAP condition (indicated with arrows). Upper band: Cx36-EGFP; lower band: Cx36-SNAP. **(B)** Confocal scans were used to confirm that V5-GBP-TurboID is directed to Cx36 whorls. Inset: 5 µm. **(C)** A BioID screen identifies p62 as a potential interacting protein of Cx36. Heat map of mass spec hits and heir abundance. Colocalization of p62 and Cx36-SNAP in transfected HEK293T cells. Scale: 10 µm. Magnified inset: 5 µm.

## Discussion

In the present study we demonstrate that ER retention of Cx36 results in the formation of multimembrane vesicles containing gap junction-like connexin aggregates. Our data suggest a model in which premature docking of Cx36 connexons reshapes the ER membrane into concentric whorls that are released into the cytoplasm (Figure 7). These findings are supported by site directed mutagenesis and electron micrographs revealing similar structural profiles for gap junctions and connexin whorls. This type of whorl formation appears to be an overexpression artefact that occurs because transfected cells produce enormous amounts of proteins that exceed the capacity of the ER export machinery. We even detected connexin whorls in Cx36 wildtype transfectants, which suggests that not all the synthesized proteins can leave the ER by a conventional export mechanism. This illustrates that the formation of connexin whorls is a concentration-dependent phenomenon. Each of the two Cx36 variants, Cx36/S318-ter and Cx36-SNAP, have an increased propensity to form whorls because these constructs are trapped in the ER, which ultimately promotes interactions between proximal connexons. Another important conclusion that can be drawn from our observations is that Cx36 is most likely to oligomerize in the ER, as only fully assembled connexons are capable of forming gap junctions. Hence, Cx36 follows a similar pattern as the beta group connexins Cx32 and Cx26 and forms hemichannels before reaching the Golgi apparatus (Evans et al., 1999; Falk et al., 1997; Koval et al., 1997; Musil and Goodenough, 1993).

**Figure 7.**
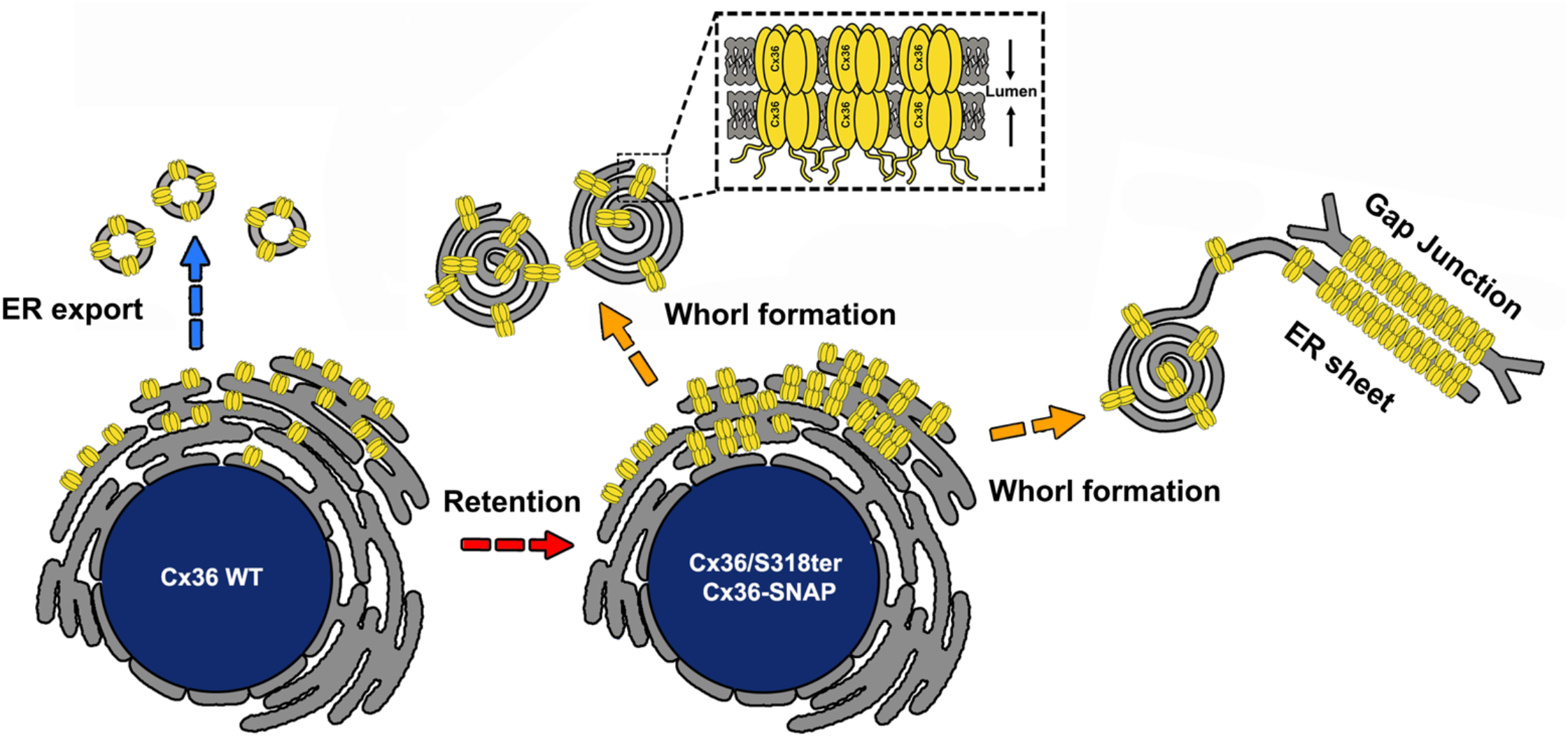
Cartoon illustrating the whorl formation mechanism. Connexin whorls are the result of ER retention. This retention promotes premature docking of connexons in the ER causing the formation of whorls.

Connexin whorls with similar features to the ones described here were reported for a cataract associated mutant of the lens connexin Cx50. In this mutant proline 88 is substituted by serine, which causes a conformational change that blocks the release of Cx50 from the ER. It has been suggested that Cx50 containing ER whorls act as light scattering particles that are detrimental for the function of the lens. To study the mechanism of whorl formation Lichtenstein et al. (Lichtenstein et al., 2009) used pulse-chase labeling and found that Cx50 whorls are formed by the addition of newly synthesized proteins to the outer layer of the vesicle. Our experiments suggest, to the contrary, that newly synthesized protein is constantly added to the inside of the whorl, whereas older Cx36 molecules form the outer layers. These differences may be ascribed to the type of tag that was used in each study. Lichtenstein et al. (2009) used a flash tag consisting of 4 cysteine whereas our experiments were conducted with the SNAP-tag. Also, it is possible that intermolecular interactions between the cytoplasmic domains of the connexin affect the orientation of adjacent ER sheets while the whorl is being formed. Hence, there are a couple of factors that could explain the differences in the whorl formation mechanism. Still, in case of both connexins, Cx50 and Cx36, it is evident that ER retention and a premature docking are the main causes of whorl formation.

Our ultrastructural examinations revealed that ER sheets were often closely connected to gap junctions and Cx36 whorls. These sheets exhibited similar structural profiles as gap junctions suggesting that they were formed by docked connexons. Comparable structures were mentioned in a review by (Segretain and Falk, 2004), who reported that rough ER sheets are tightly associated with gap junctions in HeLa cells. Their exact function is unknown, but it was proposed that these structures function as a reserve pool from which newly synthesized channels can transfer to the plasma membrane by a diffusion-based mechanism. However, it is unlikely that this is the case for the structures we have observed in this study. The proximity of these sheets and their alignment along the plasma membrane suggests that the gap junction is attracting ER-incorporated Cx36 channels through an unknown mechanism. As already pointed out before, it is possible that homophilic interactions between the cytoplasmic domains of Cx36 channels on both sides (gap junction vs. ER) lead to such an alignment. It also can’
st be excluded that certain scaffold proteins at the gap junction interact with Cx36 in ER tubules that happen to be close enough. Both scenarios we have outlined are based on the abundance Cx36 in transiently transfected cells and it is unlikely that such events occur at synapses. Nevertheless, these data essentially indicate that Cx36 gap junctions can attract intracellular connexins, thereby promoting the formation of connexin whorls at the gap junction.

ER whorls have been observed in different settings and serve as a stress response that is critical to maintain ER homeostasis (Kaiser et al., 2020; Snapp et al., 2003). In a process termed ER-phagy, whorls are released from the ER and selectively degraded by autophagosomes to ensure the timely removal of potential stressors (Yang et al., 2021). The extensive association between Cx36 and p62 implies that the degradation of connexin whorls requires the same mechanism. The p62 protein is an autophagic cargo adapter that can recruit ubiquitinated proteins into autophagosomes (Bjørkøy et al., 2005; Pankiv et al., 2007). Indeed, previous studies have shown that Cx36 is ubiquitinated by the ubiquitin ligases Lnx1 and Lnx2 (Lynn et al., 2018). These enzymes might provide the correct degradation signal for p62 to bind Cx36 and attract the expanding autophagosome. Although there is no *in vivo* evidence for the existence of connexin whorls, it is noteworthy that these structures share features with cytoprotective functions that have been reported for other systems. Connexin whorls bear a resemblance to structures such as aggresomes (Johnston et al., 1998) in the sense that they compartmentalize huge amounts of protein that might otherwise interfere with important cell functions. Moreover, our data suggest that the formation of connexin whorls functions as a mechanism to expose dense protein aggregates to a degradative pathway for cells to avoid an overabundance of protein in ER. Since overexpression systems have obvious disadvantages, it is difficult to predict whether connexin whorls are formed under *in vivo* conditions and whether they might be beneficial for cell survival. To test a potential cytoprotective function of connexin whorls we performed cell viability assays and compared the frequency of dead cells between Cx36-SNAP and Cx36-SNAP/C62 transfected HEK293T cells. These tests revealed no apparent differences in cell viability nor an upregulation of ER stress markers, which seems to exclude a cytoprotective function. However, we often observed that cells expressing larger amounts of the cysteine mutants displayed deformed nuclei and a rounded shape, which are typical indicators of apoptosis and cell detachment. It is important to point out that each of the cysteine mutant variants we have generated was expressed at a much lower level. It seems possible that those cells that expressed larger amounts of the mutant died immediately whereas cells producing lower concentrations survived. Our viability assays might have only included low expressing cells without any obvious signs of cytotoxicity, which could explain these conflicting results. The differences in the expression levels, however, may as well arise from different turnover rates. Pulse chase experiments have shown that Cx50 whorls in HeLa cells are long lived and exhibit an estimated half-life time of 67 h, which is about 20 times longer than that of gap junction incorporated Cx36 (Lichtenstein et al., 2009; Wang et al., 2015). Since ER whorls contain multiple layers of densely packed connexins, they may be less prone to degradation, which would extend their half-life time and ultimately increase the expression level of Cx36.

Our data indicate that the PDZ binding motif of Cx36 is critical for the export from the ER. The deletion of the SAYV sequence resulted a striking phenotype that was characterized by the formation of ER whorls and an aggregation of Cx36 vesicles around the nucleus. This finding apparently raises the question of whether there is a certain PDZ protein that interacts with Cx36 in the early secretory pathway to control the delivery of connexons to the gap junction. Indeed, PDZ domain containing scaffold proteins such as syntenin1 have been shown to mediate the ER export of pro-TGF-α (Fernandez-Larrea et al., 1999). A similar mechanism could apply to Cx36, which would explain the increased ER retention we observed for the Cx36/S318ter mutant lacking the PDZ binding motif. However, our data do not make a case that the PDZ ligand of Cx36 functions as a sort of ER export signal. Such export signals are often quite short and consist of 1 or 2 amino acids (Yin et al., 2017). It is possible that certain amino acids within in the sequence of the PDZ binding motif constitute a separate export signal. Previous studies have shown that a single C-terminal valine residue at position -1 (C-terminal tip) is sufficient to drive COPII dependent ER export of the Major Histocompatibility Complex, Class I, F (HLA-F) (Cho et al., 2010). Interestingly, valine is also the C-terminal amino acid of Cx36, suggesting that a similar export mechanism could be involved. The main task of ER export signals is to mediate an interaction between the cargo protein (Cx36) and specific cargo receptors, so called Sec24 proteins. These proteins recruit the cargo into COPII vesicles, which are subsequently released from the ER and transported to the Golgi (Wendeler et al., 2007). It is possible that the Cx36/318ter mutant is unable to interact with these cargo receptors because it is missing the corresponding binding motif. This would ultimately prevent the incorporation of the mutant into export vesicles resulting in ER retention and subsequent whorl formation. Surprisingly, we observed that the Cx36/318-ter mutant still formed gap junctions although a huge percentage of the connexin was trapped in the ER. This suggests that there must be an alternative mechanism to export Cx36 from the ER. Meyer et al., (2014) have previously reported that neurons such as the AII amacrine cell use different assembly mechanisms for gap junctions with different synaptic partners. One feasible explanation is a 2^nd^ export signal that might be sufficient to partially sustain the transport of the mutant. However, further studies will be required to elucidate the ER export mechanisms of Cx36.

The present study demonstrates that a premature docking of connexons functions as removal mechanism compartmentalizing Cx36 channels that are retained in the ER. Further studies will be necessary to confirm if such a mechanism operates in neurons to avoid an overabundance of connexins in the ER. As neurodegenerative diseases such as Parkinson’s or Alzheimer have been suggested to involve ER stress (Lindholm et al., 2006), connexin whorls could represent a mechanism to support ER homoeostasis in stressed cells.

## Acknowledgement

We would like to thank Ya-Ping Lin and Nikki Brantley for the excellent technical assistance. We are thankful for the support of the live cell imaging experiments by the Center for Advanced Microscopy, Department of Integrative Biology & Pharmacology at McGovern Medical School, UTHealth. We’
sd like to thank Li Li from the Clinical and Translational Proteomics Service Center for the analysis of the mass spectrometry data. S.T was supported by the *Deutsche Forschungsgemeinschaft* (DFG) (TE 1459/1-1, Walter Benjamin stipend). K.D. acknowledges funding from the DFG (RTG 1885/2 Molecular Basis of Sensory Biology) and the European Union under the action of ERA-NET NEURON (JTC2020: Rethealthsi), financed by the German Federal Ministry of Education and Research (BMBF, 01EW2107). Additional support was provided by NIH grant R01EY012857 (JO) and core grant P30EY028102. L.B. & V.L. were funded by the *Deutsche Forschungsgemeinschaft* (DFG iBiOs (No. PI 405/14-1), SFB 944 Z-Project).

## Notes

### Competing Interest Statement

The authors have declared no competing interest.

